# Single-molecule view of coordination in a multi-functional DNA polymerase

**DOI:** 10.1101/2020.08.12.248633

**Authors:** Raymond F. Pauszek, Rajan Lamichhane, Arishma Rajkarnikar Singh, David P. Millar

## Abstract

Replication and repair of genomic DNA requires the action of multiple enzymatic functions that must be coordinated in order to ensure efficient and accurate product formation. Here we have used single-molecule FRET microscopy to investigate the physical basis of functional coordination in DNA polymerase I (Pol I) from *E. coli*, a key enzyme involved in lagging-strand replication and base excision repair. Pol I contains active sites for template-directed DNA polymerization and 5’ flap processing in separate domains. We show that a DNA substrate can spontaneously transfer between polymerase (*pol*) and 5’ nuclease (*5’ nuc*) domains during a single encounter with Pol I. Additionally, we show that the flexibly tethered *5’ nuc* domain adopts different positions within Pol I-DNA complexes, depending on the nature of the DNA substrate. Our results reveal the structural dynamics that underlie functional coordination in Pol I and are likely relevant to other multi-functional DNA polymerases.

## Introduction

DNA polymerases from many organisms need to coordinate multiple enzymatic activities to achieve accurate and efficient replication and repair of DNA, while avoiding the formation of mutagenic or unstable DNA intermediates (***Reha-Krantz, 2010; Bębenek and Ziuzia-Graczyk, 2018***). DNA polymerase I (Pol I), a key enzyme involved in DNA replication and repair in *E. coli* (***Makiela-Dzbenska et al., 2009; Imai et al., 2007; Patel et al., 2001***), contains three distinct enzymatic activities in a single 928 residue polypeptide: a DNA-dependent 5’-3’ polymerase (*pol*), a proofreading 3’-5’ exonuclease (*exo*) and a 5’ nuclease (*5’ nuc*) (***Kelley and Joyce, 1983; Setlow and Kornberg, 1972***). The *pol* and *exo* activities are contained in separate domains, which together comprise the main body of the enzyme, whereas the *5’ nuc* activity is located in an independent domain that is tethered to the body by an unstructured 16 amino acid (aa) peptide linker. The *5’ nuc* domain is related to the flap endonuclease (FEN) family of structure-specific DNA nucleases (***Harrington and Lieber, 1994***).

Pol I plays an important role in lagging strand DNA replication in *E. coli* (***Balakrishnan and Bambara, 2013; Okazaki et al., 1971***). During this complex process, short RNA primers anneal to the lagging strand and are extended by DNA primase, producing fused RNA-DNA strands (Okazaki fragments). The nascent DNA portion is subsequently extended by Pol I, until the growing strand encounters another Okazaki fragment lying downstream, displacing the 5’ end and forming an RNA flap. The *5’ nuc* activity of Pol I then cleaves the RNA flap, generating a nicked DNA substrate that is subsequently sealed by a DNA ligase (***Figure 1***A). The same processing steps are also performed by Pol I during DNA base excision repair, in which case the displaced strand is composed of DNA (***Imai et al., 2007***).

**Figure 1.**
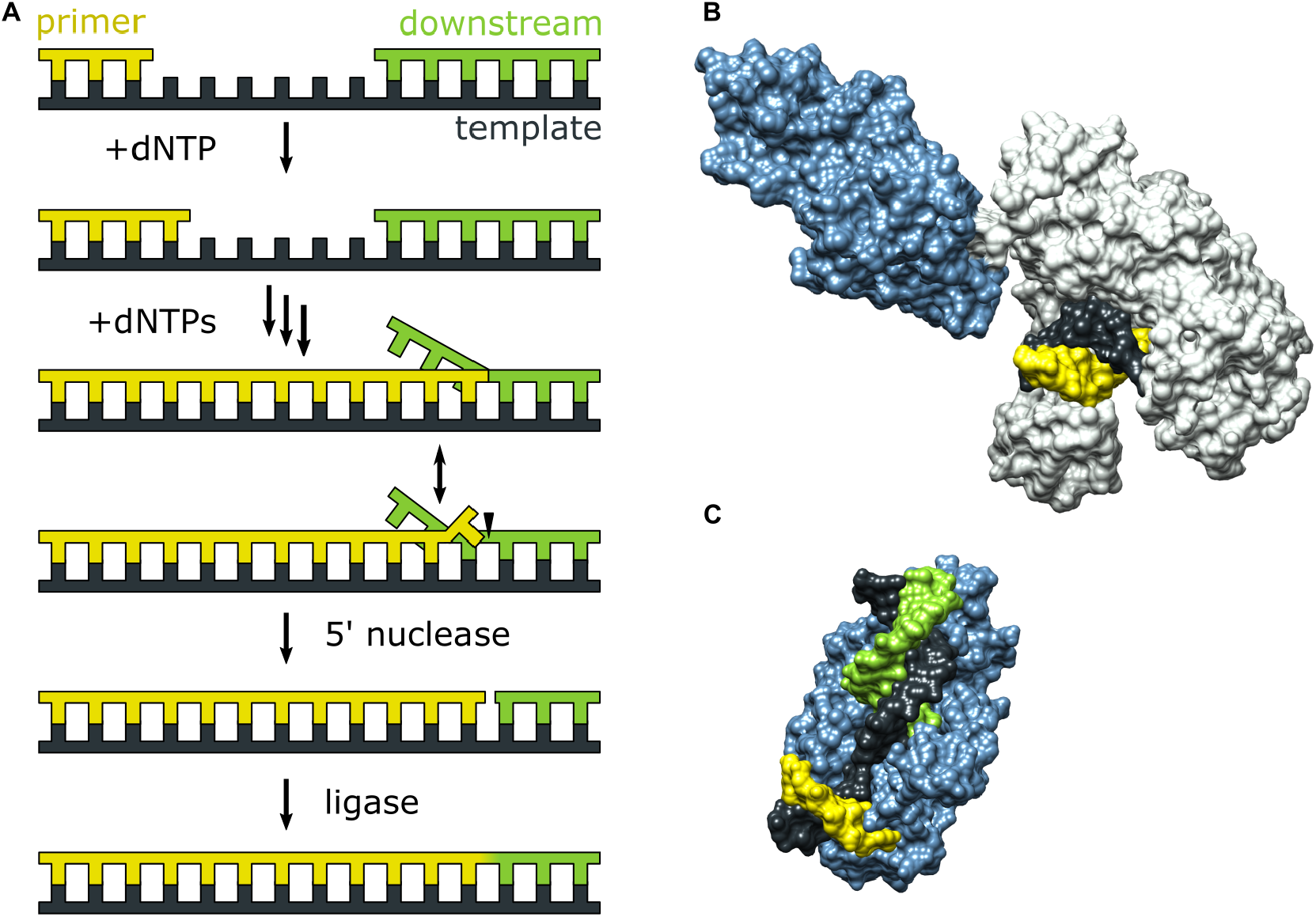
Activities of Pol I and three-dimensional structures of Pol I homologs. (A) Processes catalyzed by Pol I during lagging strand DNA replication or base excision repair. Agrowing primer strand is extended through repeated cycles of nucleotide incorporation, resulting in displacement of a downstream strand (RNA in the case of Okazaki fragment maturation or DNA in the case of base excision repair). The resulting substrate rearranges to a double flap structure that is cleaved by the 5’ nuclease activity of Pol I (site indicated by a half arrow head). The Anal ligation step is performed by a DNA ligase (not shown). (B) Crystal structure of the Pol I homolog *Taq* polymerase with DNA primer/template bound in the *pol* domain (PDB ID: 1TAU). The polymerase core is shown in gray and the *5’ nuc* domain is shown in light blue. (C) Crystal structure of human FEN1 bound to a DNA substrate (PDB: 3Q8M). In both B & C, the DNA strands are colored as in A.

During either Okazaki fragment processing or base excision repair, Pol I must achieve an appropriate balance between the *pol* and *5’ nuc* activities in order to generate a nicked duplex product that can subsequently be sealed by a DNA ligase (***Mortusewicz et al., 2006***). Excessive *pol* activity can generate long downstream 5’ flaps that are difficult to cleave (***Reha-Krantz, 2010***), while excessive *5’ nuc* activity can result in extended regions of single-stranded DNA that are prone to breakage (***Zheng et al., 2005***). Either outcome is deleterious to genomic integrity and cellular survival. A previous biochemical study suggests that 5’ flap cleavage is performed by the same Pol I molecule that extended the primer (***Xu et al., 2000***), indicating a high degree of functional coordination. However, the structural and physical basis for this coordination is unknown, since crystal structures of Pol I engaged with DNA substrates via the *pol* or *5’ nuc* domains have not been reported to date. However, the structure of the Pol I homolog *Taq* polymerase bound to DNA (***Eom et al., 1996***) reveals that the primer 3’ terminus is located within the palm region of the *pol* domain, while the *5’ nuc* domain is extended away from the enzyme core (***Figure 1***B). Moreover, the structure of human FEN1, which is homologous to the *5’ nuc* domain of Pol I, in complex with a DNA substrate provides a model for how the *5’ nuc* domain of Pol I engages a DNA substrate (***Tsutakawa et al., 2011***). The structure shows a single unpaired base at the primer 3’ terminus inserted into a pocket of FEN1 and a region of extended contacts between the protein and the sugar-phosphate backbones of both strands of the downstream DNA (***Figure 1***C). These structures of homologous enzymes suggest that the *pol* and *5’ nuc* binding modes of Pol I must differ significantly, raising questions about how the enzyme switches from one mode of activity to another.

Theoretical modelling of the transition from *pol* to *5’ nuc* activity in Pol I, starting from the extended enzyme conformation shown in ***Figure 1***B, suggests that the *5’ nuc* domain flips by ~180° and adopts a position above the fingers and thumb subdomains within the enzyme core (***Xie and Sayers, 2011***), facilitating interactions with the primer terminal base and a downstream DNA flap. Additionally, it is likely that the primer terminal base must detach from the *pol* domain in order to insert into the *5’ nuc* domain, as seen in the FEN1-DNA structure (***Figure 1***C). However, these hypothetical movements of the DNA substrate and *5’ nuc* domain have not been observed experimentally.

Here, we use single-molecule Förster resonance energy transfer (smFRET) microscopy to investigate the physical basis of functional coordination in Pol I and to probe large scale conformational changes of the enzyme-DNA complex. The smFRET method has been previously applied to Klenow fragment (KF), which is derived from Pol I by removal of the *5’ nuc* domain. These studies have monitored DNA synthesis by KF (***Christian et al., 2009***), detected nucleotide-induced conformational transitions within KF (***Santoso et al., 2010; Berezhna et al., 2012; Hohlbein et al., 2013***), monitored a DNA primer switching between the *pol* and *exo* sites of KF (***Lamichhane et al., 2013***) and established a structural model of a DNA substrate bound to KF (***Craggs et al., 2019***). However, smFRET has not yet been applied to full length Pol I, or any other DNA polymerase containing a *5’ nuc* activity.

In this study, we have developed a smFRET system to identify separate subpopulations of DNA engaging the *pol* and *5’ nuc* domains of Pol I, revealing how the fractional populations and dwell times of each species vary according to the nature of the DNA substrate. We have also implemented a complementary smFRET system to probe the location of the flexibly tethered *5’ nuc* domain, revealing that this domain undergoes a large positional shift in order to interact with a downstream DNA strand. Importantly, using both smFRET systems, we demonstrate that DNA substrates can switch between the *pol* and *5’ nuc* domains during a single encounter with Pol I. Based on kinetic analysis of relevant transitions, we propose a physical model for intramolecular transfer of DNA between the *pol* and *5’ nuc* domains. Altogether, the information from the smFRET experiments reported here provides new insights into the physical mechanisms and enzyme conformational changes that underlie functional coordination in Pol I and are likely relevant to other multifunctional DNA polymerases with spatially separated active sites.

## Results

### Experimental Design

We designed a series of model DNA substrates containing elements expected to engage the *5’ nuc* domain of Pol I. The substrates are shown in schematic form in ***Figure 2*** and the complete sequences of the constituent oligonucleotides are listed in ***Appendix 1***. One substrate contains a single-stranded 5’ flap on the downstream strand, referred to as downstream flap DNA (***Figure 2***A). Another substrate contains the downstream flap and a single unpaired 3’ terminal base on the primer strand (referred to as double flap DNA, ***Figure 2***B). A third substrate can potentially exist as a mixture of downstream flap and double flap forms (referred to as mixed flap DNA, ***Figure 2***C). We also examined DNA substrates containing a nick or gap of various sizes (***Figure 2***D) or lacking a downstream strand entirely (***Figure 2***E). In all cases, the template contains a dT_10_ spacer and biotin group at the 3’ end for surface attachment. Individual encounters between Pol I, present in solution, and the surface-immobilized DNA substrates were visualized by smFRET microscopy. Two different FRET labelling schemes were employed, as described in the following sections.

**Figure 2.**
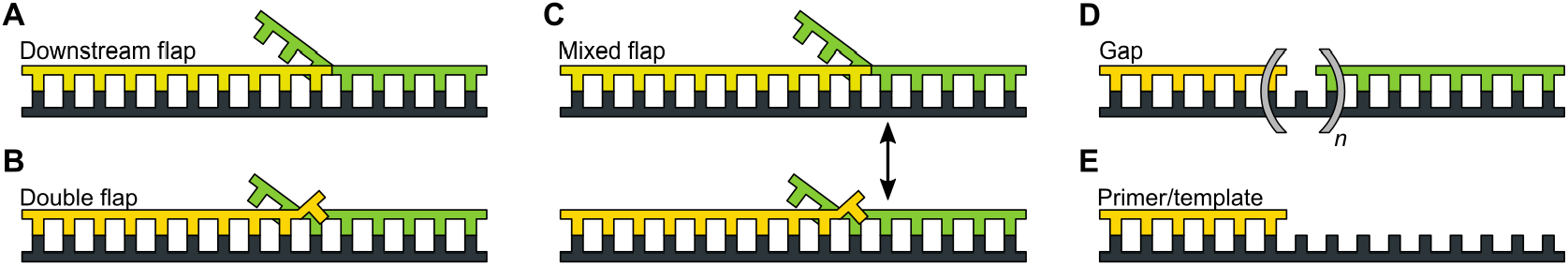
DNA substrates used in this study. (A) Substrate containing a 5’ flap on the downstream strand (designated downstream flap DNA). (B) Substrate containing the same 5’ flap as in A, plus a single unpaired base at the primer 3’ terminus (designated double flap DNA). Because of the base sequences of the strands (***Appendix 1***), the structures shown in A and B are “locked in”. (C) Substrate that can exist as a mixture of the structures shown in A and B (designated mixed flap DNA). (D) Substrates containing a nick (*n*=0) or gaps of various size (*n*=1-4). (E). Primer/template substrate.

### Movement of DNA between *pol* and *5’ nuc* domains of Pol I

The first FRET system was designed to probe the location of the DNA substrates relative to the enzyme core. A base located within the primer strand was labelled with an Alexa Fluor 488 (A488) FRET donor and Pol I was labelled at position 550 in the thumb region with a complementary Alexa Fluor 594 (A594) FRET acceptor (***Figure 3***A and ***Appendix 1-Table 1***). To achieve site-specific labelling of Pol I, the two native cysteines were removed via C262S and C907S mutations, and a single cysteine was introduced at the desired labelling site via a K550C mutation. The Pol I construct also contained D424A and D116A mutations to eliminate 3-5’ exonuclease and 5’ nuclease activities, respectively (***Derbyshire et al., 1991; Xu et al., 2001***). Pol I was expressed and purified as described in the Methods section and each step of the purification process was monitored by PAGE (***Appendix 2***).

**Figure 3.**
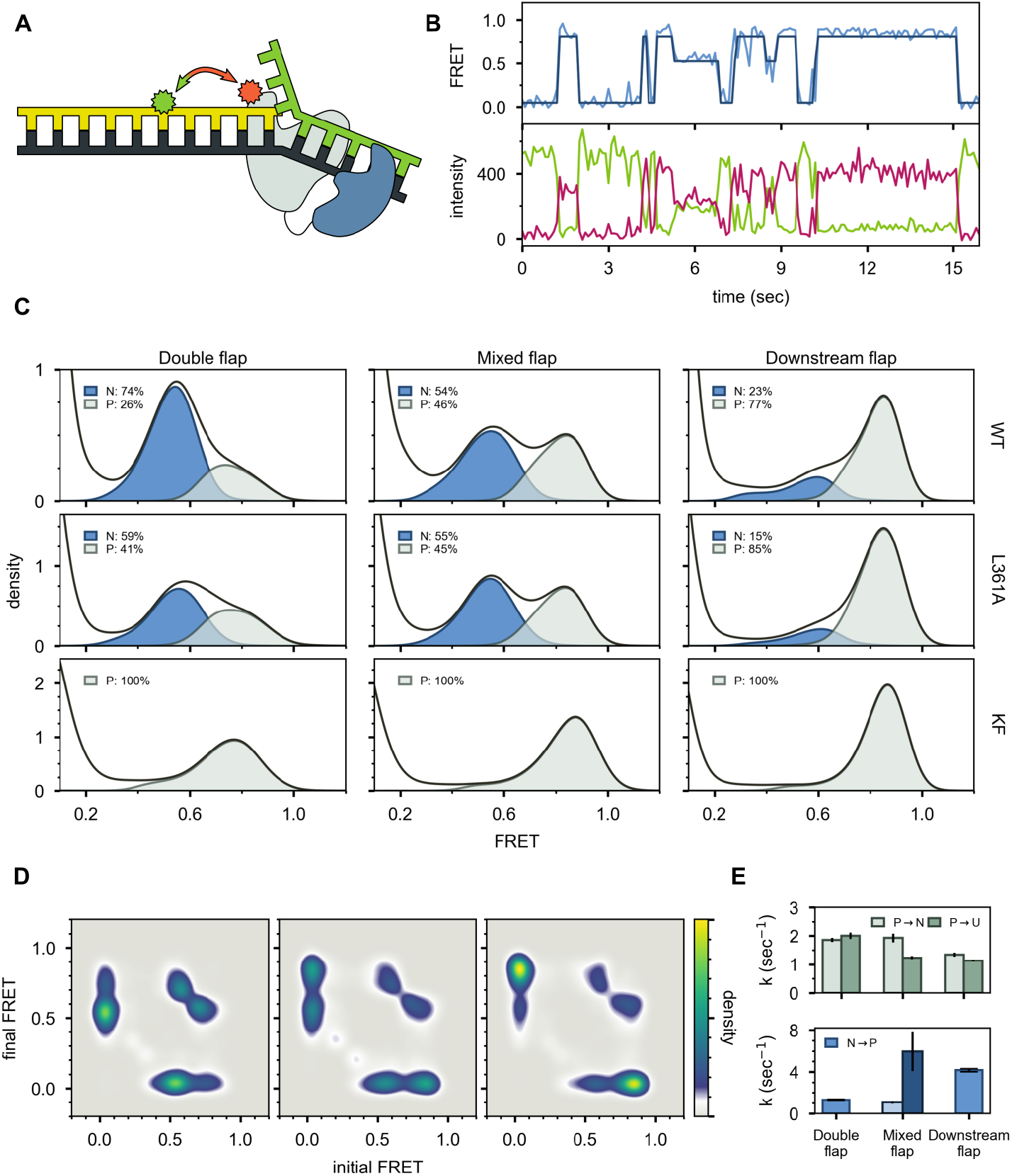
Probing the location of DNA substrates within Pol I. (A) Schematic representation of the donor (green, attached to primer strand) and acceptor (orange, attached to thumb region of Pol I) labelling sites. Pol I is depicted in cartoon form, with the core coloured grey and the *5’ nuc* domain coloured blue. (B) Representative smFRET trajectory (blue) and donor (green) and acceptor (magenta) emission trajectories, for Pol I interacting with mixed flap substrate. The bold line is the idealized state path determined from Hidden Markov modelling. (C) Composite FRET efficiency histograms for states P and N, compiled during global Hidden Markov analysis, for various combinations of DNA substrate and protein. The proteins, from top to bottom, are WT Pol I, Pol I L361A and KF L361A. The corresponding populations of states P and N are indicated. (D) Transition density plots for Pol I interacting with flap-containing DNA substrates. From left to right: double flap DNA, mixed flap DNA and downstream flap DNA. (E) Rate constants for intramolecular transfer of DNA substrates from *pol* domain to *5’ nuc* domain (P→N), compared with overall dissociation rates from the *pol* domain (P→U, where U denotes unbound). (F) Rate constants for intramolecular transfer of DNA substrates from *5’ nuc* domain to *pol* domain (N→P). The mixed flap DNA exhibits two transfer rates (***Figure 3-Figure Supplement 1***), both of which are shown. **Figure 3-Figure supplement 1.** Dwell time histograms for Pol I interacting with flap-containing substrates **Figure 3-Figure supplement 2.** Histogram of emission intensities resulting from direct excitation of A594 in Pol I bound to downstream flap DNA.

A representative set of donor, acceptor and FRET efficiency trajectories depicting a series of encounters between Pol I and immobilized DNA (mixed flap in this example) is shown in ***Figure 3***B. During each encounter, the donor intensity abruptly drops, and an acceptor signal appears at the same instant, reflecting binding of Pol I to the DNA, while at a later time point the acceptor signal disappears and the donor signal increases correspondingly, reflecting dissociation of Pol I from the DNA. The anti-correlation of the donor and acceptor signals confirms that the intensity changes are due to FRET. During the binding periods, the FRET efficiency alternates between two distinct levels at ~0.8 or ~0.6 efficiency. Multiple FRET trajectories for each DNA substrate interacting with Pol I were analysed globally using a Hidden Markov model, confirming that two distinct states are sufficient to account for all data sets. The resulting composite FRET efficiency histograms for each state are shown in ***Figure 3***C. Note that each histogram is accumulated during the global HMM analysis, not by Gaussian fitting of final envelopes, ensuring clean separation of the two states and accurate quantification of the state populations.

A state at 0.8 FRET efficiency was previously observed for DNA primer/templates interacting with KF (same donor and acceptor labelling sites as here) and was attributed to DNA engaging the *pol* domain (***Lamichhane et al., 2013***). Since KF is identical to the main core of Pol I, the 0.8 FRET state observed here is also attributed to DNA engaging the *pol* domain within Pol I (state P). To test whether the 0.6 FRET state arises from DNA binding to the *exo* domain of Pol I, we introduced a L361A mutation, which is known to disrupt DNA binding at the *exo* domain of KF (***Lamichhane et al., 2013; Lam et al., 1998***). The L361A mutation has little effect on the 0.6 FRET population (compare top and middle rows in ***Figure 3***C), indicating that this state does not arise from binding of DNA to the *exo* domain. We also examined KF, which lacks the *5’ nuc* domain entirely. The KF construct was also labelled at position 550 with A594 and contained an L361A mutation. Notably, the 0.6 FRET population is not observed with KF (***Figure 3***C, bottom row). Together, these observations indicate that the 0.6 FRET state arises from DNA engaging the *5’ nuc* domain of Pol I (state N).

The lower FRET efficiency of state N could be due to movement of the upstream duplex (where the donor is located) as the primer terminus detaches from the *pol* domain and moves to the *5’ nuc* domain, tilting of the protein helix to which the acceptor is attached, or changes in local environment of donor and/or acceptor that alter the Förster radius. Photophysical control experiments indicate that the Förster radius in state N is similar to that in state P (***Appendix 3***), indicating that the lower FRET efficiency of state N is due to physical movement of the DNA substrate and/or thumb subdomain. However, since the thumb subdomain is a rigid structural element within Pol I and KF, it is likely that the FRET change is actually due to movement of the DNA substrate. The change in FRET efficiency from 0.8 to 0.6 corresponds to a lengthening of the donor-acceptor distance of ~7 Å (***Appendix 3***).

State N is most highly populated for the double flap substrate (***Figure 3***C, top left), consistent with the known preference of the *5’ nuc* activity of Pol I for double flap structures (***Xu et al., 2001, 2000***). In contrast, state N is least populated with the downstream flap substrate (***Figure 3***C, top right). The FRET histogram for the mixed flap substrate reveals an intermediate population of state N (***Figure 3***C, top centre), consistent with this substrate existing as a mixture of double flap and downstream flap forms.

Two-dimensional transition density plots (TDPs) (***McKinney et al., 2006***) were constructed from multiple FRET trajectories to reveal the connectivity of the various FRET states (Fig. 3D). The two peaks evident on the y-axis reflect binding of Pol I to the immobilized DNA, using either the *pol* or *5’ nuc* domains. Dwell time analysis reveals that the on-rates for binding via either domain are similar, and in the range expected for a diffusion-controlled binding process (***Appendix 4-Table 1***), indicating that both domains are equally accessible. The peaks evident on the x-axis reflect the corresponding dissociation transitions. Dwell time analysis shows that the double flap substrate exhibits the fastest dissociation from the *pol* domain and the slowest dissociation from the *5’ nuc* domain (***Appendix 4-Table 1***). The equilibrium dissociation constants calculated from the kinetic rate constants are also listed in ***Appendix 4-Table 1***.

These results imply that Pol I can engage DNA via one domain (*pol* or *5’ nuc*), dissociate into bulk solution, and subsequently rebind using the other domain. An example of such an event sequence (dissociate from state N and rebind in state P) is evident in the representative smFRET trajectory shown in ***Figure 3***B (middle portion, from ~6s to ~8 s). Dissociation and rebinding provide one pathway for transfer of DNA substrates between *pol* and *5’ nuc* domains (intermolecular transfer). In addition, we observe frequent transitions between states P and N that do not show a measurable pause in a zero-FRET state (an example is also shown in ***Figure 3***B, between ~8 s and ~10s). Such transitions give rise to prominent cross peaks between 0.6 and 0.8 FRET states in the TDPs (***Figure 3***D). The rate constants for these direct transitions from state P to N (intramolecular transfer) were determined by dwell time analysis for each DNA substrate (***Figure 3-Figure Supplement 1*** and ***Appendix 4-Table 2***). Interestingly, the transfer rates forthe double flap and downstream flap substrates are similar to the corresponding rates at which Pol I bound via the *pol* domain dissociates into bulk solution (***Figure 3***E, upper panel).

There are two models that account for the direct transitions between states P and N, and the correspondence between the transfer rates and the dissociation rates. First, Pol I bound to DNA via the *pol* domain could spontaneously dissociate and be re-captured by the same DNA molecule via the *5’ nuc* domain before escaping into bulk solution. Alternatively, two Pol I molecules could bind to a single DNA substrate, one engaging the DNA via the *pol* domain and the other engaging the DNA via the *5’ nuc* domain. Dissociation of the first Pol I molecule would give rise to a transition from 0.8 to 0.6 FRET efficiency. To distinguish these possibilities, we evaluated the stoichiometry of Pol I on each of the DNA substrates. To do so, we excited the A594 acceptor directly and recorded the resulting A594 emission over time, under the same conditions used for the smFRET experiments. A histogram of A594 emission intensity, compiled from >100 individual Pol I molecules interacting with immobilized DNA (downstream flap in this example), reveals a peak at ~350 a.u. (***Figure 3-Figure Supplement 2***), corresponding to a single Pol I molecule bound to DNA (the peak at zero intensity is due to periods in which Pol I is not bound to the DNA). Notably, there is no indication of a peak or shoulder at ~700 a.u., corresponding to two bound Pol I molecules. Similar results were obtained for the other DNA substrates (not shown). We conclude that a single Pol I molecule is bound to DNA under the conditions of our experiments, ruling out the second scenario described above. Based on counting of the relevant transitions for double flap DNA, 64% of exit events from the *pol* domain result in transfer to the *5’ nuc* domain, while the remaining 36% result in escape into the bulk solution (***Appendix 4-Table 3***).

Interestingly, the *pol* to *5’ nuc* transfer rate for the mixed flap substrate matches that for the double flap substrate (***Figure 3***E, upper panel). As noted above, the mixed flap substrate can potentially exist as a mixture of double flap and downstream flap forms. The kinetic results in ***Figure 3***E suggest that the rate of transfer from the *pol* domain to the *5’ nuc* domain is governed by the double flap form, whereas overall dissociation from the *pol* domain reflects both double flap and downstream flap forms.

Dwell time analysis reveals significant variations in the rates of intramolecular transfer from state N to state P among the various DNA substrates (***Figure 3***E, lower panel & ***Appendix 4-Table 2***). The downstream flap substrate exhibits the fastest transfer. In contrast, transfer of the double flap substrate is much slower, showing the importance of the unpaired primer base for stable engagement with the *5’ nuc* domain. The structure of FEN1 (homologous to the *5’ nuc* domain of Pol I) with DNA reveals that the unpaired primer base is sequestered in a binding pocket, making contacts with a network of surrounding protein residues (***Tsutakawa et al., 2011***). A similar arrangement in Pol I would account for the prolonged residence time of DNA in state N and the slow return to state P. Interestingly, the dwell time histogram for the mixed flap substrate does not fit well to a single exponential function and instead requires a biexponential function for best fit (***Figure 3-Figure Supplement 1***). Moreover, the rate constants for the fast and slow kinetic phases match the transfer rates observed for the downstream flap or double flap substrates, respectively (***Figure 3***E, lower panel). This observation is consistent with the expectation that the mixed flap substrate exists as a mixture of downstream flap and double flap forms, and further shows that the two subpopulations can be resolved kinetically.

### Role of downstream strand and 5’ flap for binding to the *5’ nuc* site

To investigate whether the presence of a downstream strand is required to engage the *5’ nuc* domain, we examined a DNA substrate containing only primer and template strands (***Figure 2***E and ***Appendix 1-Table 1***). This substrate exhibits a single FRET state at ~0.8 efficiency, corresponding to state P, while interacting with Pol I (***Figure 4***A & B, bottom panels). Likewise, no cross peaks are observed in the TDP (***Figure 4***C). State N is absent in this case, showing that the downstream DNA must be in duplex form to engage the *5’ nuc* domain and promote movement of the upstream DNA out of the *pol* domain.

**Figure 4.**
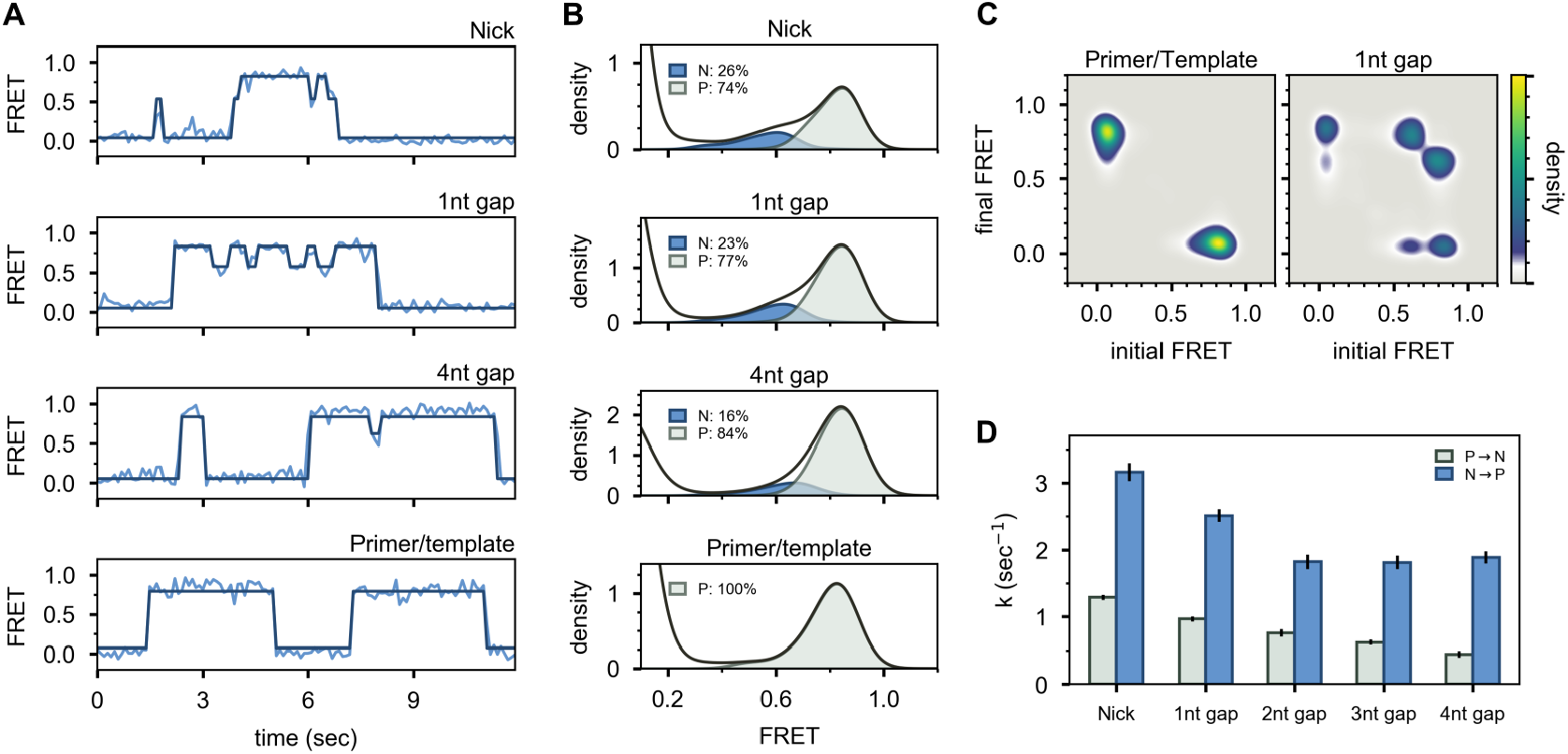
Interaction of Pol I with primer/template DNA or DNA substrates containing a nick or gap. (A) Representative smFRET trajectories for DNA substrates interacting with Pol I, as indicated. Schematic representations of DNA substrates are shown in ***Figure 2***. Bold lines are idealized state paths determined from Hidden Markov modelling. (B) Composite FRET efficiency histograms for states P and N, for various DNA substrates interacting with Pol I, as indicated. The corresponding populations of states P and N are indicated. (C) Transition density plots for Pol I interacting with various DNA substrates, as indicated. (D) Rate constants for intramolecular transfer of various DNA substrates between *pol* domain and *5’ nuc* domain (P→N, grey) or between *5’ nuc* domain and *pol* domain (N→P, blue). **Figure 4-Figure supplement 1.** Histograms and TDPs of Pol I L361A and KF interacting with 1nt gap. **Figure 4-Figure supplement 2.** Dwell time histograms for Pol interacting with nick-or gap-containing substrates

To test whether the presence of a 5’ flap on the downstream strand is also required to engage the *5’ nuc* domain, we examined DNA substrates containing a nick or single-stranded gaps of various sizes (***Figure 2***D and ***Appendix 1-Table 1***). These substrates reveal reversible transitions between ~0.6 and ~0.8 FRET states when interacting with Pol I (***Figure 4***A, upper three panels), similar to the substrates containing 5’ flaps. Moreover, the hallmarks of the 0.6 FRET state are indicative of DNA engaging the *5’ nuc* domain (state N, ***Figure 4-Figure Supplement 1***). The populations of state N for the nicked and single-nucleotide gap substrates (26% and 23%, respectively, ***Figure 4***B) are similar to the downstream flap substrate (23%, ***Figure 3***C), indicating that the presence of a downstream strand is sufficient to engage the *5’ nuc* domain, regardless of whether that strand contains a 5’ flap or not. Moreover, the *5’ nuc* domain can engage a downstream strand even when it is separated from the primer strand by a 4nt gap (***Figure 4***A & B), which is likely a consequence of the flexible 16 aa linker tethering the *5’ nuc* domain to the body of the enzyme. The flexibility of the single-stranded gap region could also facilitate docking of the *5’ nuc* domain with the downstream DNA strand. The presence of prominent cross peaks in TDPs shows that a gapped DNA substrate can transfer reversibly between *pol* and *5’ nuc* domains during a single encounter with Pol I (***Figure 4***C), while dwell time analysis (***Figure 4-Figure Supplement 2***) shows that transfer from the *pol* domain to the *5’ nuc* domain becomes progressively slower as the gap size increases (***Figure 4***D and ***Appendix 4-Table 4***). Likewise, return of DNA from the *5’ nuc* domain to the *pol* domain is also slower for the larger gap sizes (***Figure 4***D and ***Appendix 4-Table 4***). The rate constants for binding or dissociation of primer/template, nicked and gapped DNA substrates are listed in ***Appendix 4-Table 5***.

### Movement of the *5’ nuc* domain during docking with downstream DNA

The second FRET system is designed to probe the proximity of the *5’ nuc* domain to the downstream portion of the DNA substrates. In this case, an A488 donor was attached to a cysteine residue introduced at position 213 in the *5’ nuc* domain of Pol I (via aT213C mutation) and an A594 acceptor was attached to a base in the downstream portion of the template strand (***Figure 5A & Appendix 1-Table 2***). The DNA substrates were otherwise analogous to those used in scheme 1. The A594 acceptor was placed in the template, rather than downstream strand, to avoid any possible steric interference with binding of Pol I. The Pol I construct also contained C262S, C907S, D424A and D116A mutations, as described above. A representative set of donor, acceptor and FRET efficiency trajectories for the mixed flap DNA substrate is shown in ***Figure 5***B. For this labelling scheme, there is no signal from either donor or acceptor during the periods when Pol I is not present on the DNA. Upon binding of Pol I, both donor and acceptor signals appear simultaneously and their relative magnitudes reflect the FRET efficiencyduringthe encounter. Upon dissociation of Pol I, both signals disappear at the same instant (***Figure 5***B).

**Figure 5.**
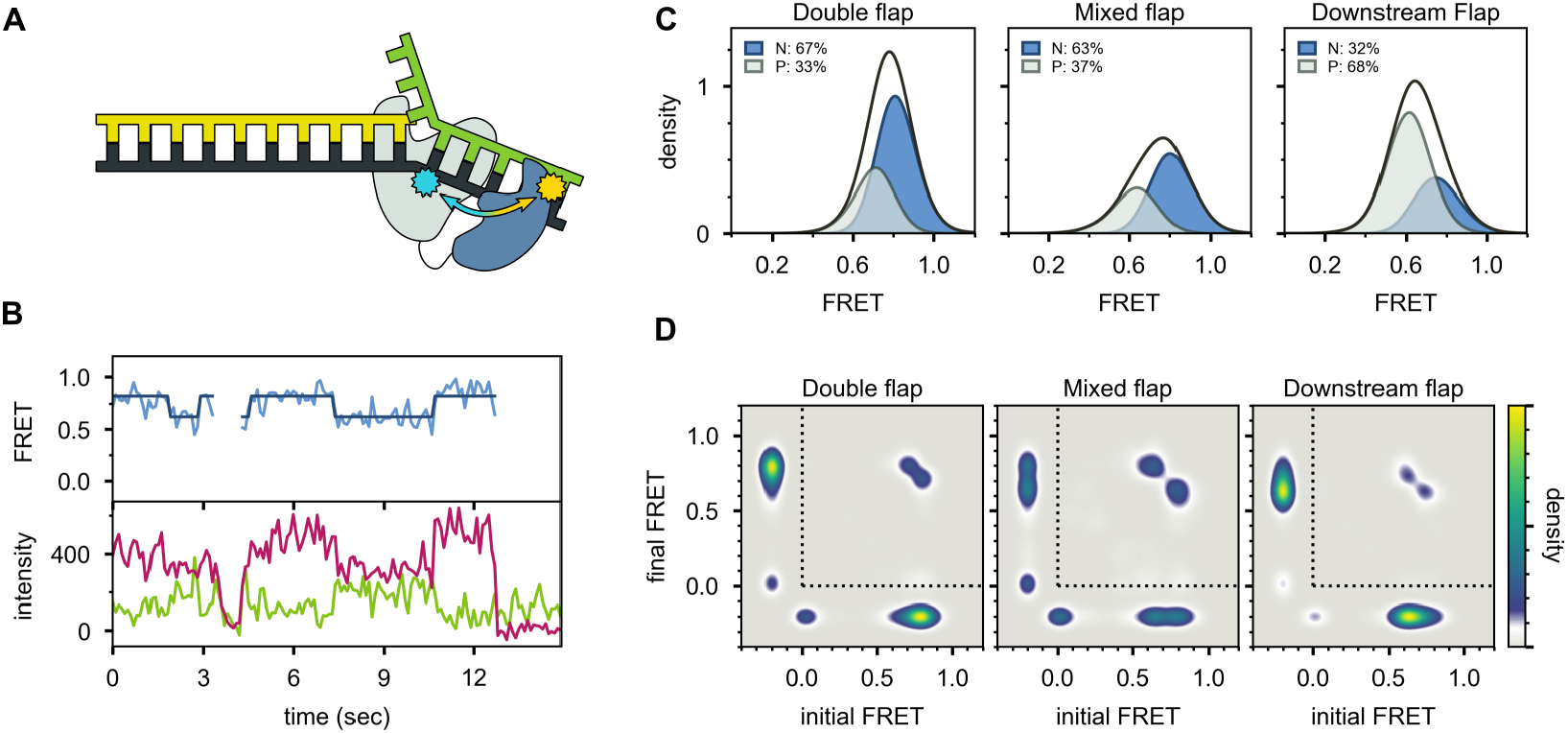
Probing the location of the *5’ nuc* domain within Pol I-DNA complexes. (A) Schematic representation of donor (yellow, attached to *5’ nuc* domain) and acceptor (cyan, attached to downstream template strand) labelling sites. (B) Representative set of donor, acceptor and FRET trajectories for mixed flap DNA substrate. (C) FRET efficiency histograms compiled from multiple trajectories. (D) Transition density plots compiled from same data sets in (C). Since the FRET efficiency is not defined during periods when Pol I is not bound to DNA, the FRET efficiency is set to −0.2. The quadrant enclosed by the dotted lines corresponds to periods during which Pol I is bound to DNA.

Multiple FRET trajectories for each DNA substrate (containing 5’ flap) interacting with Pol I were analysed globally using a Hidden Markov model, showing that two distinct states were sufficient to account for all data sets. The resulting FRET efficiency histograms for each state are centred at ~0.8 and ~0.6 efficiency (***Figure 5***C). Notably, the fractional populations of the high-FRET and mid-FRET peaks are similar to the fractional populations of states N and P, respectively, observed with the first labelling scheme (compare ***Figure 5***C and ***Figure 3***C). This is true across all three substrates containing 5’ flaps, which partition differently between states N and P. Accordingly, the high-FRET and mid-FRET species observed with the present labelling scheme are also assigned to states N and P, respectively. The high FRET efficiency of state N indicates that the donor and acceptor sites are relatively close in space (donor-acceptor distance ~40Å), confirming that state N arises from engagement of the *5’ nuc* domain with the downstream DNA. The lower FRET efficiency of state P indicates that the *5’ nuc* domain is somewhat further from the downstream DNA when the primer 3’ terminus occupies the *pol* domain (donor-acceptor distance ~49 Å). Direct transitions between the mid-FRET and high-FRET states are observed in individual FRET trajectories (***Figure 5***B) and in composite TDPs (***Figure 5***D), indicating that Pol I can switch between states P and N without dissociation. This conclusion is consistent with the observations from the first labelling scheme (***Figure 3***D).

In striking contrast, the primer/template substrate mostly populates a single FRET state with a much lower efficiency of ~0.3 (***Figure 6***), indicating that the *5’ nuc* domain is distant from the downstream template (donor-acceptor distance ~60Å). We designate this state P’(see Discussion). The FRET histogram also reveals a barely detectable shoulder at ~0.6 efficiency, corresponding to state P (***Figure 6***C). There are very few transitions between states P’ and P, as indicated by the absence of cross peaks in the TDP (***Figure 6***D). Overall, these observations are consistent with the first labelling scheme, which indicates that the downstream DNA must be present in duplex form in order to engage the *5’ nuc* domain (***Figure 4***B).

**Figure 6.**
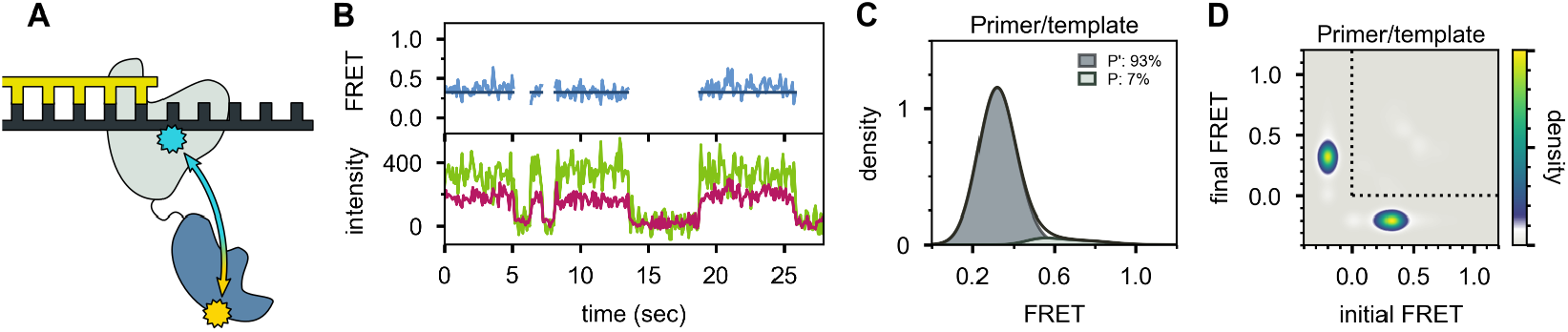
Location of the *5’ nuc* domain within Pol I bound to primer/template substrate. (A) Schematic representation of donor (yellow) and acceptor (cyan) labelling sites. (B) Representative set of donor, acceptor and FRET trajectories. (C) FRET efficiency histograms compiled from multiple trajectories. (D) Transition density plot compiled from the same data sets as in C. Same presentation as ***Figure 5***D.

## Discussion

The *5’ nuc* domain of Pol I fulfils a key function during DNA replication and repair, cleaving 5’ flaps that arise from strand displacement synthesis (***Figure 1***A). The physical and structural mechanisms that underlie the coordination between the *pol* and *5’ nuc* activities of Pol I are not well understood, owing to technical challenges in studying this enzyme. The *5’ nuc* domain is tethered to the enzyme core by an unstructured 16 aa peptide linker, which may allow this domain to adopt a range of positions. This intrinsic flexibility has likely impeded efforts to crystallize Pol I and determine a high-resolution crystal structure. More fundamentally, the potential ability of the *5’ nuc* domain to exchange between different positions within a Pol I-DNA complex could playa role in orchestrating the transition from one mode of enzymatic activity to another.

Here, we have developed two complementary smFRET systems to visualize Pol I spontaneously exchanging between different DNA binding modes and to probe the location of the *5’ nuc* domain in each case. Using the first labelling scheme, we have resolved two distinct FRET states and shown that they correspond to DNA engaging the pol domain (state P) or engaging the *5’ nuc* domain (state N). While we have previously observed the *pol* binding mode in analogous smFRET studies of KF (***Lamichhane et al., 2013***), the present spectroscopic observations of the *5’ nuc* binding mode in full length Pol I have not been reported before. Using the second labelling scheme, we have confirmed that state N arises from docking of the *5’ nuc* domain with the downstream DNA. Notably, the fractional populations of states P and N determined with each labelling scheme are in agreement (***Figure 3***C & ***Figure 5***C), establishing a consistent description of the Pol I-DNA complexes under study. In contrast, if a downstream strand is not present, the *5’ nuc* domain adopts a different position, extended way from the DNA substrate (***Figure 6***), underscoring the mobility of this independent and flexibly tethered protein domain.

Although structures of full-length Pol I with DNA substrates are not available, our spectroscopic data on states P and N are consistent with details revealed in the structures of homologous enzymes bound to DNA substrates. The crystal structure of *Taq* polymerase with a DNA primer/template engaging the *pol* domain shows that the primer 3’ terminal base is located in a cleft within the palm region (***Eom et al. (1996)***, ***Figure 1***B). In contrast, the structure of FEN1 (homologous to the *5’ nuc* domain of Pol I) in complex with a DNA substrate reveals an unpaired 3’ terminal base located in a pocket surrounded by a cluster of protein residues (***Tsutakawa et al. (2011)***, ***Figure 1***C). In the context of Pol I, these two structures suggest the primer terminal base detaches from the *pol* domain and inserts into the *5’ nuc* domain, which is consistent with our observation that the DNA duplex undergoes a ~7 Å displacement between states P and N. Moreover, the observation that the primer terminal base is unpaired in the FEN1 structure is consistent with our spectroscopic observations that state N is most highly populated with the double-flap DNA substrate (***Figure 3***C & ***Figure 5***C). The FEN1 structure also shows a region of extended contacts between the protein and the sugar-phosphate backbones of both strands of the downstream DNA (***Tsutakawa et al., 2011***), consistent with our observations that the downstream DNA must be in duplex form to engage the *5’ nuc* domain (***Figure 4***B) and that a downstream strand engages the *5’ nuc* domain of Pol I even when separated from the primer terminus by a 4nt gap (***Figure 4***B).

An important finding from our smFRET study is that DNA substrates can transfer reversibly between *pol* and *5’ nuc* domains during a single encounter with Pol I. These intramolecular transitions are readily observable in individual FRET trajectories (***Figure 3***B & ***Figure 5***B) and in composite transition density plots (***Figure 3***D & ***Figure 5***D), obtained using either labelling scheme. Our results explain previous biochemical observations indicating that the same Pol I molecule that extends the primer terminus can also cleave the resulting downstream flap (***Xu et al., 2000***). An intramolecular transfer pathway is also employed during *pol* to *exo* switching in KF (***Lamichhane et al., 2013; Joyce, 1989***). Most importantly, our results provide insights into the physical mechanisms that facilitate the transfer of DNA from the *pol* domain to the *5’ nuc* domain in Pol I. The dwell time analysis shows that the transfer rate matches the rate of dissociation from the *pol* domain into bulk solution for the double flap and downstream flap substrates (***Figure 3***E). These results suggest a model for intramolecular transfer, whereby Pol I bound via the *pol* domain spontaneously detaches from the DNA and immediately reengages the same DNA molecule via the *5’ nuc* domain before escaping into bulk solution.

In fact, the *5’ nuc* domain appears to be positioned appropriately to act in this fashion. Based on the second FRET system, the donor-acceptor distance in state P (~49 Å) is in a similar range to state N (~40 Å), suggesting that the *5’ nuc* domain is in relatively close proximity to the downstream DNA when the primer 3’ terminus is located in the *pol* domain. This positioning increases the likelihood that the *5’ nuc* domain will capture the same DNA substrate after Pol I dissociates. This is consistent with our observation that 64% of all transitions out of the *pol* domain result in direct transfer to the *5’ nuc* domain (***Appendix 4-Table 3***). In contrast, if a downstream DNA strand is not present, the *5’ nuc* domain adopts a more distant position, with a donor-acceptor distance of ~60 Å (***Figure 6***B & C). We do not observe any direct transitions between *pol* and *5’ nuc* domains under these conditions (***Figure 4***C and ***Figure 6***D).

Taken together, the results from the two FRET systems suggest that complexes of Pol I with DNA can adopt three distinct configurations, summarized schematically in ***Figure 7***. Complex P’ is formed exclusively with a primer/template substrate: the primer 3’ terminus is bound in the *pol* domain and the *5’ nuc* domain is extended away from the enzyme core. The prime symbol is to distinguish this species from complex P, which forms with DNA substrates containing a downstream strand: the primer 3’ terminus is still located in the *pol* domain, but the *5’ nuc* domain is in proximity to the downstream strand. Complexes P and P’ are indistinguishable using the first labelling scheme, but are clearly resolved with the second scheme, highlighting the importance of utilizing multiple donor and acceptor sites to detect all species present. In complex N, the *5’ nuc* domain is in closest proximity to the downstream strand and the primer terminus has shifted from the *pol* domain to the *5’ nuc* domain. Our results show that complexes P and N can freely interconvert, with the distribution of the two species being determined by the nature of the DNA substrate. Complex P is the dominant species for the downstream flap and 1nt gap substrates, with transient excursions to complex N. In contrast, the double flap substrate is biased towards complex N, consistent with the known substrate preference of the *5’ nuc* activity of Pol I (***Xu et al., 2001, 2000***). Theoretical modelling also suggests that the *5’ nuc* domain can adopt different positions within a Pol I-DNA complex (***Xie and Sayers, 2011***).

**Figure 7.**
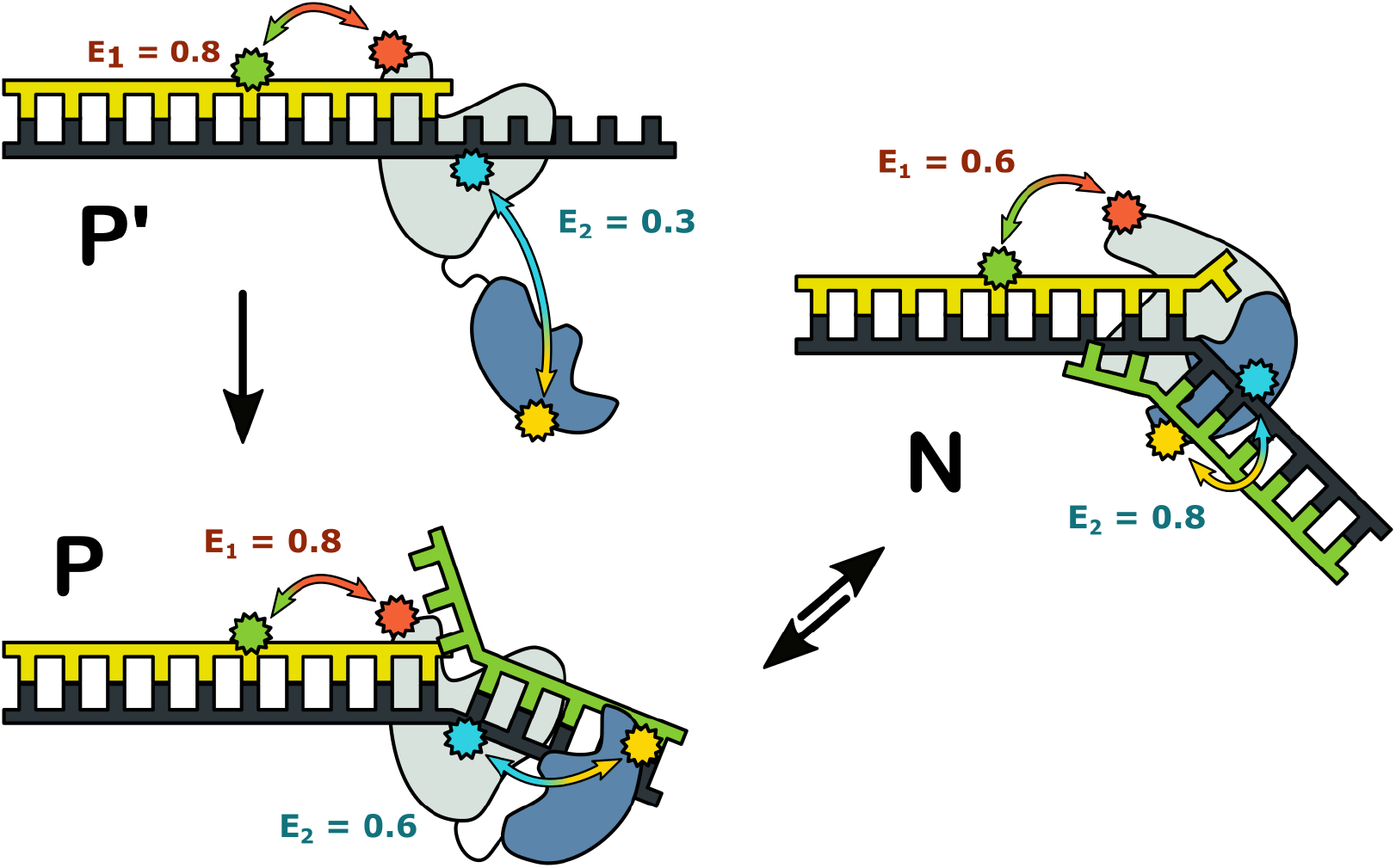
Fig. 7: Possible configurations of Pol I-DNA complexes. (A) Complex P’. In the absence of a nearby downstream strand, the primer 3’ terminus resides in the *pol* domain and the *5’ nuc* domain is extended away from the enzyme core. The donor and acceptor probes for the first labelling scheme are green and magenta, respectively, while the donor and acceptor for the second labelling scheme are yellow and blue. The FRET efficiencies for the first and second labelling schemes are denoted E_1_ and E_2_, respectively. (B) Complex P. The primer 3’ terminus resides in the *pol* domain and the *5’ nuc* domain is located in proximity to a nearby downstream strand. Other details as in A. (C) Complex N. The *5’ nuc* domain is docked with the downstream strand and the DNA primer 3’ terminus resides within the same domain. Other details as in A.

The complexes in ***Figure 7*** likely correspond to snapshots during strand displacement synthesis and the transition from *pol* to *5’ nuc* activity (***Figure 1***A). Complex P’ corresponds to an early stage, when the nascent primer strand is distant from a downstream strand. Complex P corresponds to a later stage, in which the growing primer has displaced the downstream strand but the primer 3’ terminus is still located in the *pol* domain. Complex N likely represents the next step in the pathway, in which the primer 3’ terminus has moved out of the *pol* domain and the *5’ nuc* domain is poised to cleave the scissile phosphodiester within the downstream strand, although the actual cleavage step is blocked here by a D116A mutation. Future smFRET studies with active Pol I should provide further insights into the temporal order of events and the associated enzyme and DNA conformational changes along the pathway.

DNA polymerases from many organisms also possess distinct enzymatic activities that must be carefully coordinated to ensure accurate and efficient DNA replication and repair. In DNA Pol III, the major replicative polymerase in *E. coli*, the *pol* and *exo* active sites are located in separate protein subunits within a multi-protein holoenzyme complex (***McHenry, 2011; Oakley, 2019***). A similar situation prevails in eukaryotic DNA polymerases (***Burgers and Kunkel, 2017; Raia et al., 2019***). Moreover, in eukaryotes 5’ flap cleavage is performed by a separate enzyme, such as FEN1 (***Dehé and Gaillard, 2017; Stodola and Burgers, 2017***). Pol I is a relatively simple model of multi-functional DNA polymerases because it contains three distinct activities within a single polypeptide and does not require accessory proteins for proper function. Here we have shown that DNA substrates can transfer reversibly between *pol* and *5’ nuc* domains during a single encounter with Pol I. Moreover, our results show how the flexibly tethered *5’ nuc* domain adjusts it position to facilitate transfer of DNA from the *pol* to *5’ nuc* domains. Based on these observations, we have proposed a physical mechanism for transfer and functional coordination. Intramolecular transfer of the DNA substrate, combined with protein conformational changes, orchestrates the transition from one mode of activity to another, without the need to dismantle and reassemble the enzyme-DNA complex. This paradigm for functional coordination is likely relevant to more complex multi-functional DNA polymerase holoenzyme complexes, in which the various active sites are also widely separated in space, albeit within separate protein subunits.

## Methods and Materials

### Oligonucleotides

All DNA oligonucleotides were purchased from IDT in purified form and used as delivered. Oligonucleotides containing an amino-modified dT were labelled with either Alexa Fluor 488 or Alexa Fluor 594 NHS ester (ThermoFisher Scientific) according to the manufacturer’s protocol. Template strands contained a biotin group at the 3’ end for immobilization on neutravidin-coated microscope slides. All oligonucleotide sequences and modifications are listed in ***Appendix 1-Table 1 & Appendix 1-Table 2***.

### Expression of Pol I derivatives

A Pol I expression vector was generated from the plasmid pXS67 (Yale Coli Genetic Stock Center, Strain CJ803) by site-directed mutagenesis using a QuikChange kit (Agilent) according to the manufacturer’s protocol. This construct, referred to as wild-type (WT) in the text, also carried C262S and C907S mutations to remove the two native cysteines in Pol I, a D424A mutation to suppress 3’-5’ exonuclease activity, a D116A mutation to suppress 5’ nuclease activity and a 6× histidine tag attached to the C-terminus of the protein by a Gly-Pro-Gly linker. A Pol I construct containing an additional L361A mutation was generated from the WT construct by site-directed mutagenesis using a Q5 site-directed mutagenesis kit (NEB) according to manufacturer’s protocol. KF carrying K550C, D424A and C907S mutations (and L361A, as indicated) was produced from a previously described plasmid (***Berezhna et al., 2012; Lamichhane et al., 2013***). Expression and purification of KF was carried out as previously described (***Berezhna et al., 2012; Joyce and Derbyshire, 1995***). Pol I was expressed in the same manner and purified as detailed below.

### Purification of Pol I derivatives

Cells expressing Pol I were lysed by sonication in HisTrap Buffer A(50mM Tris-HCl, pH 7.5,10mM *β*-mercaptoethanol, 10mM imidazole) supplemented with 20μM phenylmethylsulfonyl fluoride (PMSF). Cellular debris was removed by centrifugation at 7500×g for 15 minutes at 4 °C, and the clarified cell extract was loaded onto a 5-mL HisTrap HP column (GE Life Sciences) equilibrated in HisTrap Buffer A. The column was washed with 5 column volumes of HisTrap Buffer A, and protein was eluted with HisTrap Buffer B (50mM Tris-HCl, pH 7.5,10mM *β*-mercaptoethanol, 250mM imidazole). Fractions containing Pol I were then loaded onto a 5-mM HiTrap Heparin HP column (GE Life Sciences) equilibrated in Heparin BufferA(50mMTris-HCl, pH 7.5,1mM DTT). Protein was eluted over a 0-50% gradient of Heparin Buffer B (50mM Tris-HCl, pH 7.5, 1 mM DTT, 2M NaCl). All purification steps were monitored by PAGE analysis (***Appendix 2***). Fractions containing Pol I were combined and exchanged into 50mM sodium phosphate buffer, pH 7, using an Econo-Pac 10DG column (Bio-Rad) prior to labelling.

### Fluorophore labelling

Pol I or KF constructs were labelled with Alexa Fluor A594 or Alexa Fluor A488 C5 maleimide (ThermoFisher Scientific) and purified as described previously (***Berezhna et al., 2012***). Protein concentrations and labelling efficiency were calculated based on optical absorption using an extinction coefficient of *ε*_280_ = 86,180 M^−1^ cm^−1^, *ε*_590_ = 90,000 M^−1^ cm^−1^, and *ε*_495_ = 71,000 M^−1^ cm^−1^ for Pol I, A594, and A488 respectively. Labelling efficiency was typically between 65 and 100%. Purified labelled protein was concentrated using a 50kDa MWCO centrifugal filter (EMD Millipore) and stored at −80°C in buffer containing 10mM Tris-HCl, pH 7.5,1 mM EDTA, 1 mM DTT, and 50% (v/v) glycerol.

### smFRET data acquisition

smFRET data collection was performed using a custom built prism-based TIRF microscope as described previously (***Berezhna et al., 2012***). Briefly, a sample chamber was assembled with quartz slides passivated with polyethylene glycol and coated with neutravidin (***Lamichhane et al., 2010***). Biotinylated DNA substrates (20pM) were flowed into the sample chamber and allowed to equilibrate for 5 minutes. The sample chamber was washed to remove unbound substrate, and 5 nM Pol I supplemented with 1 mM propyl gallate was introduced into the chamber. All solutions were prepared in imaging buffer (50mM Tris-HCl, pH 7.5, 10mM MgCl_2_, 0.5mg mL^−1^, 1mM DTT, 50mM NaCl, and 2mM Trolox). Data were collected with 100ms integration time using a custom single-molecule data acquisition program that controlled the CCD camera. Single-molecule donor and corresponding acceptor time trajectories were extracted from movies using custom scripts written in IDL. All data acquisition software packages were downloaded from https://physics.illinois.edu/cplc/software.

### smFRET data analysis

All data analysis was carried out using custom software written in-house using Python and supporting packages. Individual donor and corresponding acceptor intensity versus time traces were corrected for background signal by subtracting the median signal in each channel after photobleaching. Acceptor intensity traces were also corrected for leakage from the donor channel, as determined previously (***Berezhna et al., 2012***). FRET efficiency trajectories were calculated as

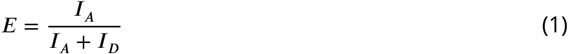

where *E* is the FRET efficiency at each time point and *I_D_* and *I_A_* are the corresponding corrected donor and acceptor intensities, respectively. Traces exhibiting anti-correlated fluctuations in the donor and acceptor intensities, constant total intensity (sum of donor and acceptor), and single-step photobleaching events were selected for further analysis.

A Hidden Markov Model (HMM) was trained globally on all selected traces for a particular protein/substrate combination simultaneously using an expectation-maximization method (***Rabiner, 1989***). For each model, the minimum number of states that adequately fit individual traces was determined by manual inspection. Once the model was trained, the Viterbi algorithm (***Rabiner, 1989***) was used to determine the most likely hidden state path for each trajectory. This labelled state path was then used to aggregate all data points belonging to a particular state in order to compile composite histograms of FRET efficiency, using a Kernel Density estimation (KDE) algorithm (scikitlearn) with a Gaussian kernel and a bandwidth of 0.04. The relative populations of distinct FRET states were directly obtained during compilation of the corresponding histograms. Transition density plots (***McKinney et al., 2006***) were constructed using a KDE (Gaussian kernel, 0.04 bandwidth), where 2D points in the training data set were defined as the median FRET efficiency in the initialand final states. Dwell-time histograms were constructed with equal bin widths across the entire data range. The optimal bin width for each histogram was estimated using the Freedman-Diaconis rule

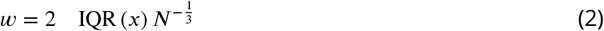

where *w* is the bin width, *x* is the array of dwell times, *N* is the number of data points in *x* and IQR is the interquartile range of the data. Kinetic rate constants and their associated uncertainties were calculated by Atting dwell-time histograms with single- or double-exponential functions (SciPy).

## Acknowledgments

We thank Edwin van der Schans for technical assistance with expression and purification of KF and Ashok Deniz for the use of the steady-state fluorimeter.

## Appendix 1

The following tables present the various DNA oligos used to make each substrate in the current study. Each row lists (from top to bottom) the primer, template, and downstream strands for each substrate. B indicates the terminal biotin group, <u>Y</u> represents the aminomodified thymine used for fluorophore labeling, while lowercase nucleotides mark mismatched (flap) regions in the primer and downstream strands. All substrates were prepared by heat-annealing a mixture of oligos at 95 °C for 5 minutes, followed by cooling slowly to room temperature on the lab bench. All mixtures contained a 3:1:3 molar ratio of primer, template, and downstream strands in order to ensure that all immobilized substrates contained all strands.

**Appendix 1 Table 1.**
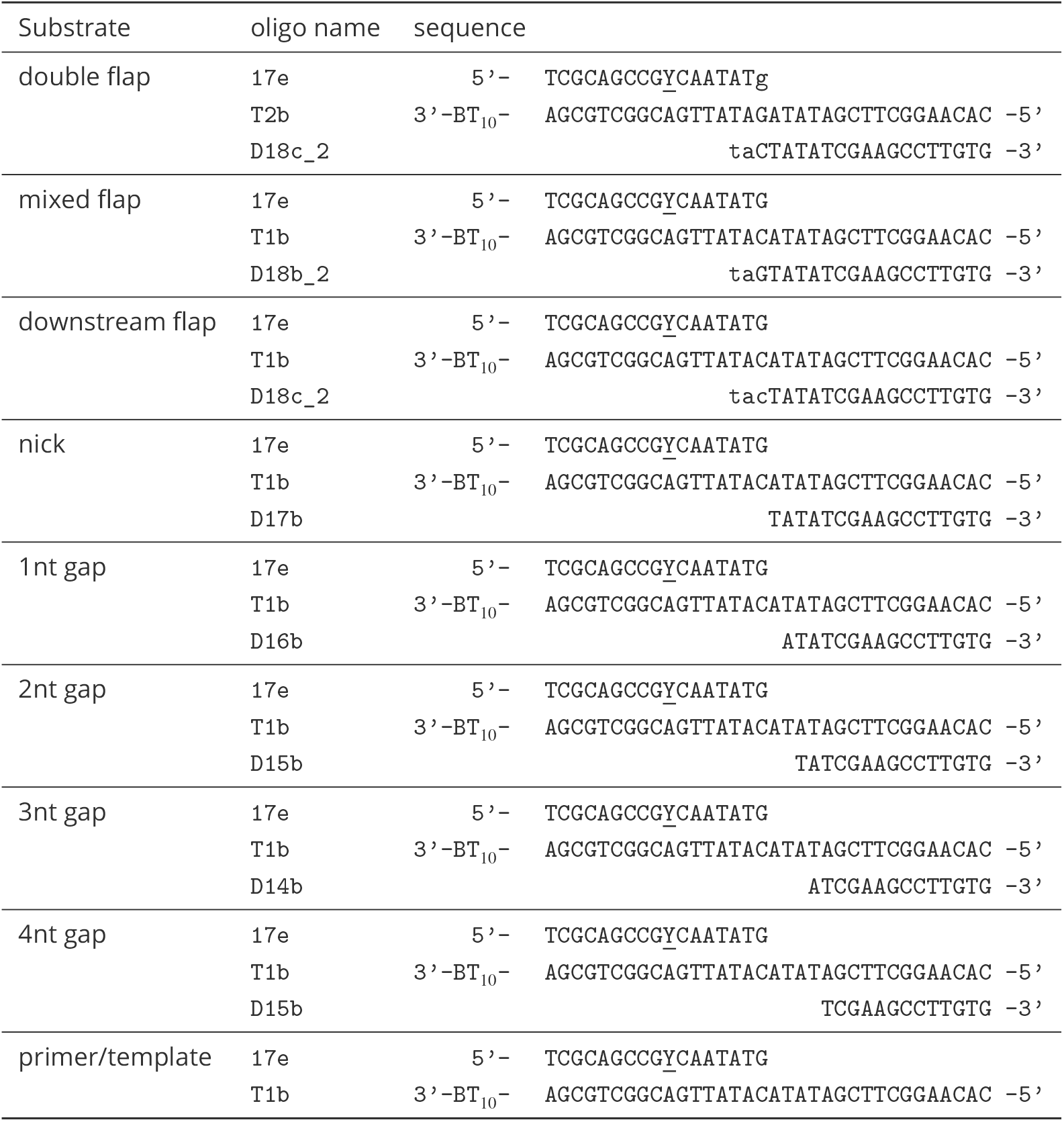
Sequences of DNA oligonucleotides used to construct substrates for FRET scheme 1.

**Appendix 1 Table 2.**
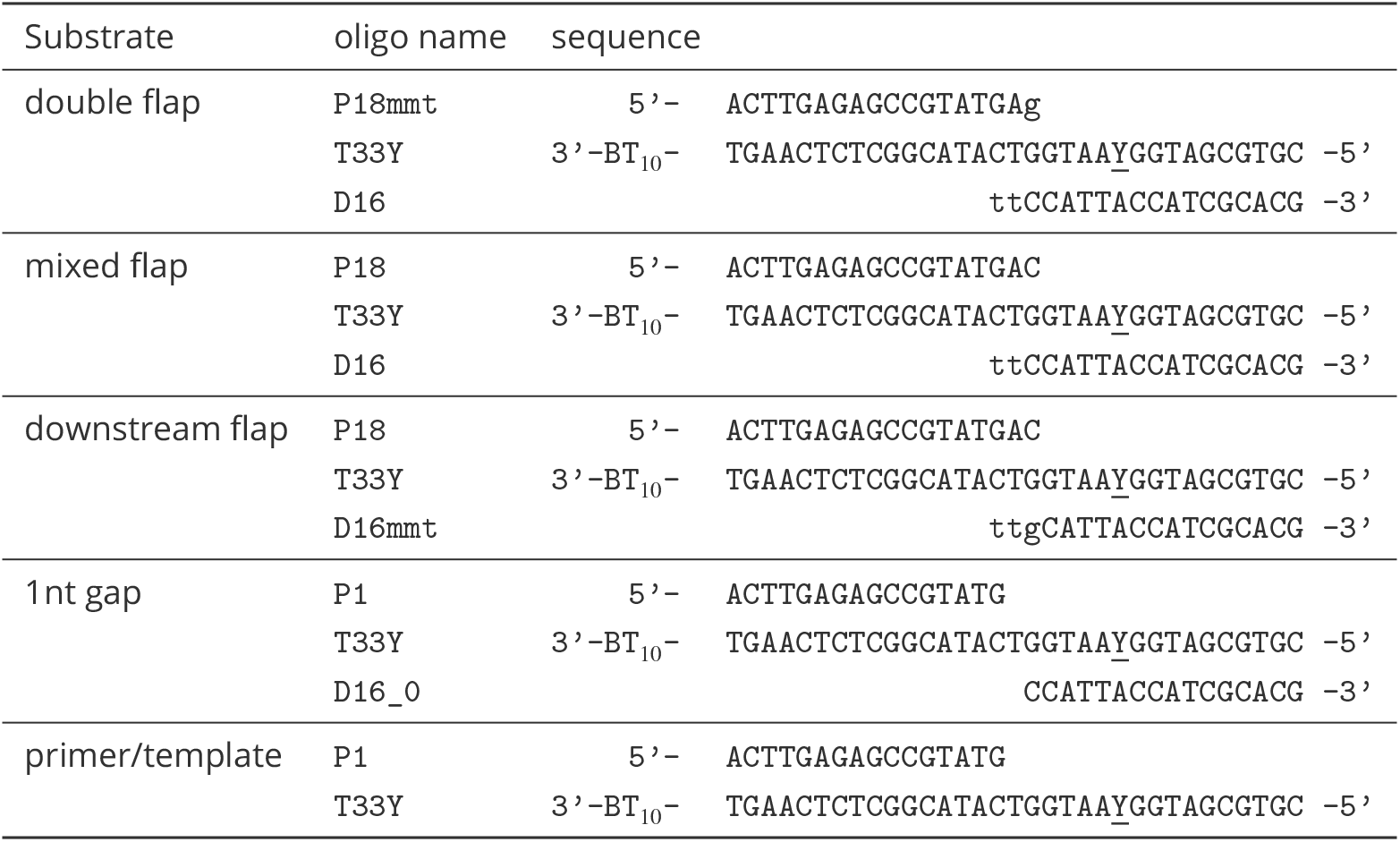
Sequences of DNA oligonucleotides used to construct substrates for FRET scheme 2.

## Appendix 2

**Appendix 2 Figure 1.**
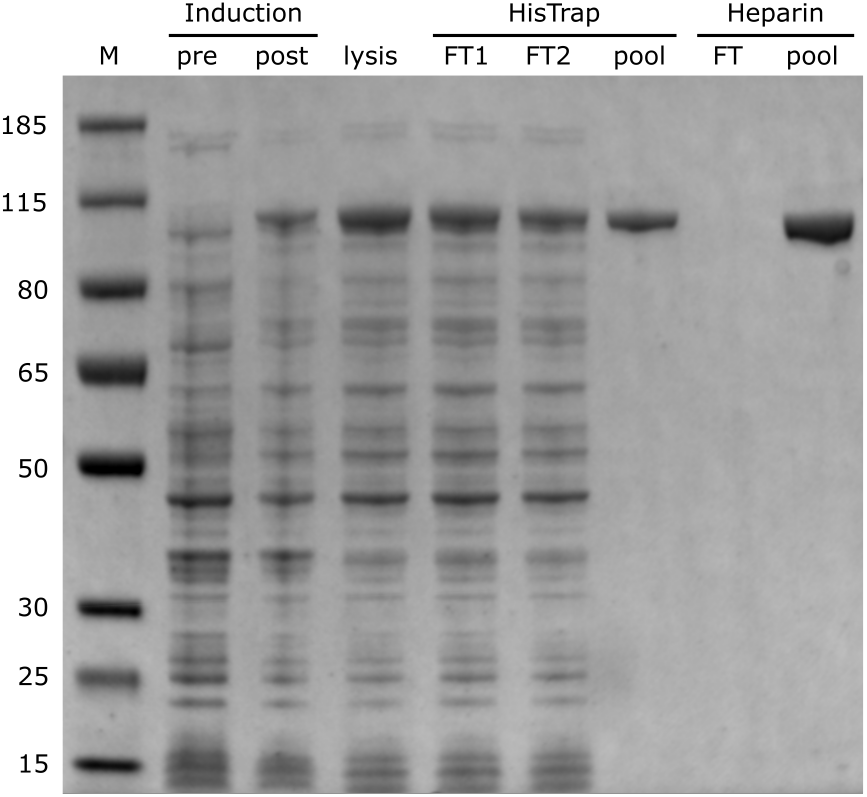
PAGE analysis of Pol I expression and purification steps. (M = size marker;FT = flow-through)

## Appendix 3

Control experiments were performed to determine whether there are any changes in spectroscopic parameters between states Pand N that could alter the Förster radius and thereby cause changes in FRET efficiency. Steady-state emission spectra and polarization anisotropy values were acquired for A488-labeled DNA substrates (***Appendix 1 – Table 1***), either alone or in the presence of saturating concentration of unlabeled Pol I. Similarly, emission spectra and anisotropy values were acquired for A594-labeled Pol I (labeled at position 550), either free in solution or in the presence of excess unlabeled DNA. The results are summarized in ***Appendix 3-Table 1***.

The primer/template exhibits a small increase in total emission intensity of A488 in the presence of unlabeled Pol I. In contrast, there is essentially no change in A488 intensity upon binding of Pol I to the double flap DNA substrate. The A488 anisotropy increases significantly in the presence of Pol I, as expected for binding of a large protein to DNA, but the final anisotropy values are similar for each bound substrate. Hence, the local rotational mobility of A488 must be similar for each bound DNA.

The emission intensity of A594 attached to Pol I shows little change upon binding of any ofthe DNA substrates, indicatingthatthe local environment of A594 is unchanged. Likewise, the anisotropy of A594 is very similar in all bound complexes, indicating that the local rotational mobility of the probe is also unchanged. The Förster radius (*R*_0_) is dependent on the donor quantum yield (*φ_D_*), the spectral overlap of donor and acceptor (*J*), the orientation factor (*κ*^2^), and other parameters, as described by the following equation

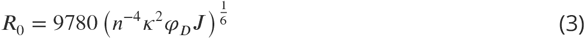

where *n* is the refractive index of the surrounding medium and *R*_0_ is in units of angstroms (***Lakowicz, 2006***). Since the donor and acceptor anisotropies are very similar in each DNA-protein complex, it is likely that the orientation factor has the same value in each case. The spectral overlap must also be similar for each complex, since we do not observe any spectral shifts in donor emission or acceptor absorbance (not shown). The one quantity that does vary to some degree among the complexes is the donor quantum yield. Assuming an intrinsic *R*_0_ value of 55.6 Å for the A488/A594 pair (***Gansen et al., 2018***), and using the normalized donor intensities in ***Appendix 3–Table 1***, we predict *R*_0_ values of 57.0 Å for the bound primer/template and 55.0 Å for the double flap substrate. Since the matched primer/template populates state P exclusively (***Figure 3***C), the *R*_0_ value for this state must also be 57.0 Å. The double flap substrate partitions between states P (26%) and N (74%), implying that the intrinsic *R*_0_ value for state N is somewhat shorter than 55.0 Å. Overall, we conclude that the Förster radii for states P and N are very similar.

The apparent donor/acceptor distance *R* corresponding to FRET efficiency *E* was calculated as follows

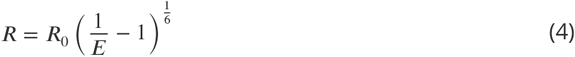

**Appendix 3 Table 1.** Steady-state fluorescence controls

| labeled macromolecule | label | ligand | normalized intensity<sup>a,b</sup> | anisotropy<sup>a</sup> |
| --- | --- | --- | --- | --- |
| double flap | A488 | none | 1.0 | 0.05 ± 0.01 |
|  |  | Pol I | 0.94 | 0.21 ± 0.05 |
| primer/template | A488 | none | 1.0 | 0.05 ± 0.01 |
|  |  | Pol I | 1.2 | 0.22 ± 0.01 |
| Pol I | A594 | none | 1.0 | 0.32 ± 0.02 |
|  |  | double flap | 1.2 | 0.34 ± 0.03 |
|  |  | primer/template | 1.1 | 0.34 ± 0.03 |

<sup>a</sup> total emission intensity normalized to the value for the free macromolecule
<sup>b</sup> A488 data obtained with excitation at 495 nm, A594 with excitation at 590 nm

## Appendix 4

**Appendix 4 Table 1.** Binding and dissociation kinetic & thermodynamic properties of Pol I interacting with flap-containing DNA substrates

| substrate | association ($10^7 \text{M}^{-1} \text{s}^{-1}$) | | dissociation ($\text{s}^{-1}$) | | $K_d$ (nM) | | |
| --- | --- | --- | --- | --- | --- | --- | --- |
| | $k_{U \rightarrow P}$ | $k_{U \rightarrow N}$ | $k_{P \rightarrow U}$ | $k_{N \rightarrow U}$ | P | N | overall |
| double flap | $5.1 \pm 0.5$ | $5.1 \pm 0.4$ | $2.0 \pm 0.1$ | $2.0 \pm 0.1$ | $40 \pm 5$ | $40 \pm 3$ | $20 \pm 2$ |
| mixed flap | $4.8 \pm 0.4$ | $5.5 \pm 0.6$ | $1.2 \pm 0.1$ | $2.7 \pm 0.2$ | $26 \pm 2$ | $48 \pm 6$ | $17 \pm 1$ |
| downstream flap | $5.9 \pm 0.3$ | $7.0 \pm 0.8$ | $1.14 \pm 0.03$ | $3.6 \pm 0.1$ | $19 \pm 1$ | $51 \pm 6$ | $14 \pm 1$ |

$U$ denotes unbound DNA.

**Appendix 4 Table 2.**
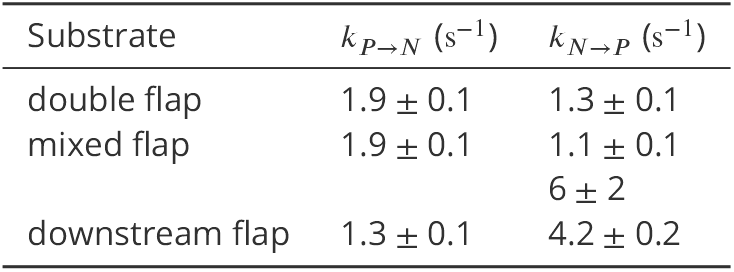
Domain-switching rate constants for Pol I interacting with flap-containing substrates

**Appendix 4 Table 3.** Statistical frequencies of transitions originating from the *pol* domain.

| substrate | $P \rightarrow N$ | $P \rightarrow U$ |
| --- | --- | --- |
| double flap | 64% | 36% |
| mixed flap | 42% | 58% |
| downstream flap | 34% | 66% |

**Appendix 4 Table 4.** Domain-switching rate constants for Pol I interacting with nick- or gap-containing substrates

| substrate | $k_{P \rightarrow N} (\text{s}^{-1})$ | $k_{N \rightarrow P} (\text{s}^{-1})$ |
| --- | --- | --- |
| nick | $1.5 \pm 0.1$ | $3.2 \pm 0.2$ |
| 1nt gap | $0.94 \pm 0.03$ | $2.5 \pm 0.4$ |
| 2nt gap | $0.76 \pm 0.05$ | $1.8 \pm 0.1$ |
| 3nt gap | $0.62 \pm 0.04$ | $1.8 \pm 0.1$ |
| 4nt gap | $0.44 \pm 0.05$ | $2.0 \pm 0.1$ |

**Appendix 4 Table 5.** Binding and dissociation kinetic & thermodynamic properties of Pol I interacting with various DNA substrates

| substrate | association ($10^7 \text{M}^{-1} \text{s}^{-1}$) | | dissociation ($\text{s}^{-1}$) | | $K_d$ (nM) | | |
| --- | --- | --- | --- | --- | --- | --- | --- |
| | $k_{U \rightarrow P}$ | $k_{U \rightarrow N}$ | $k_{P \rightarrow U}$ | $k_{N \rightarrow U}$ | P | N | overall |
| nick | $4.6 \pm 0.2$ | $5.6 \pm 0.3$ | $1.11 \pm 0.04$ | $3.5 \pm 0.1$ | $24 \pm 2$ | $62 \pm 4$ | $17 \pm 1$ |
| 1nt gap | $5.2 \pm 0.4$ | $5.6 \pm 0.8$ | $0.65 \pm 0.03$ | $2.1 \pm 0.1$ | $13 \pm 1$ | $39 \pm 6$ | $10 \pm 1$ |
| 2nt gap | $5.0 \pm 0.4$ | $6.6 \pm 0.9$ | $0.46 \pm 0.04$ | $2.4 \pm 0.2$ | $9 \pm 1$ | $37 \pm 6$ | $7 \pm 1$ |
| 3nt gap | $6.6 \pm 0.3$ | $9 \pm 1$ | $0.52 \pm 0.03$ | $1.8 \pm 0.1$ | $8 \pm 1$ | $20 \pm 2$ | $6 \pm 1$ |
| 4nt gap | $6.3 \pm 0.7$ | $13 \pm 2$ | $0.33 \pm 0.04$ | $2.4 \pm 0.2$ | $5 \pm 1$ | $19 \pm 3$ | $4 \pm 1$ |
| primer/template | $5.8 \pm 0.2$ | | $0.85 \pm 0.02$ | | | | $15 \pm 1$ |

---

**Figure 3-Figure supplement 1.**
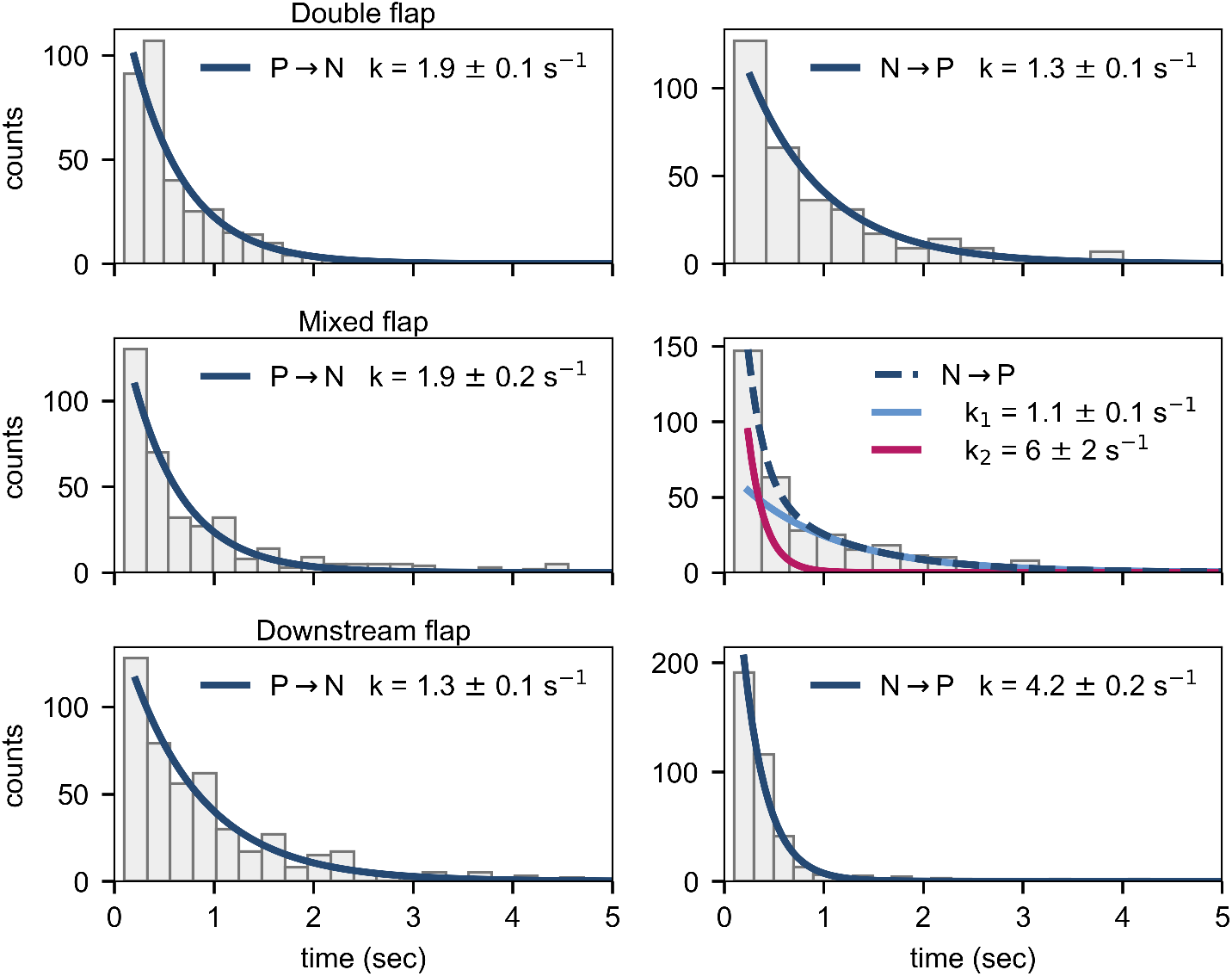
Dwell time histograms for domain-switching transitions of WT Pol I interacting with flap-containing DNA substrates. The solid lines are best fits to a single exponential function, with the rate constants indicated. The *N → P* transition for mixed flap substrate requires a biexponential function for best fit (dashed line). Individual rate constants are shown.

**Figure 3-Figure supplement 2.**
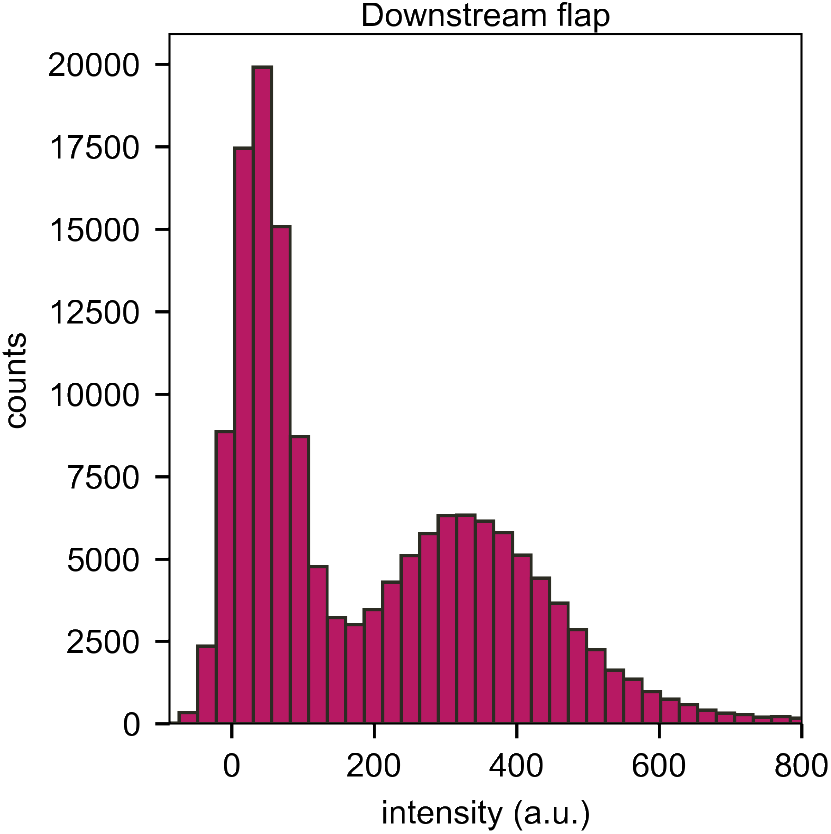
Histogram of emission intensities resulting from direct excitation of A594 in Pol I bound to downstream flap DNA. The single peak centered at ∼350 a.u. indicates that only one enzyme molecule binds to the DNA. The peak at ∼0 a.u. corresponds to periods during which Pol I is not bound to the DNA.

**Figure 4-Figure supplement 1.**
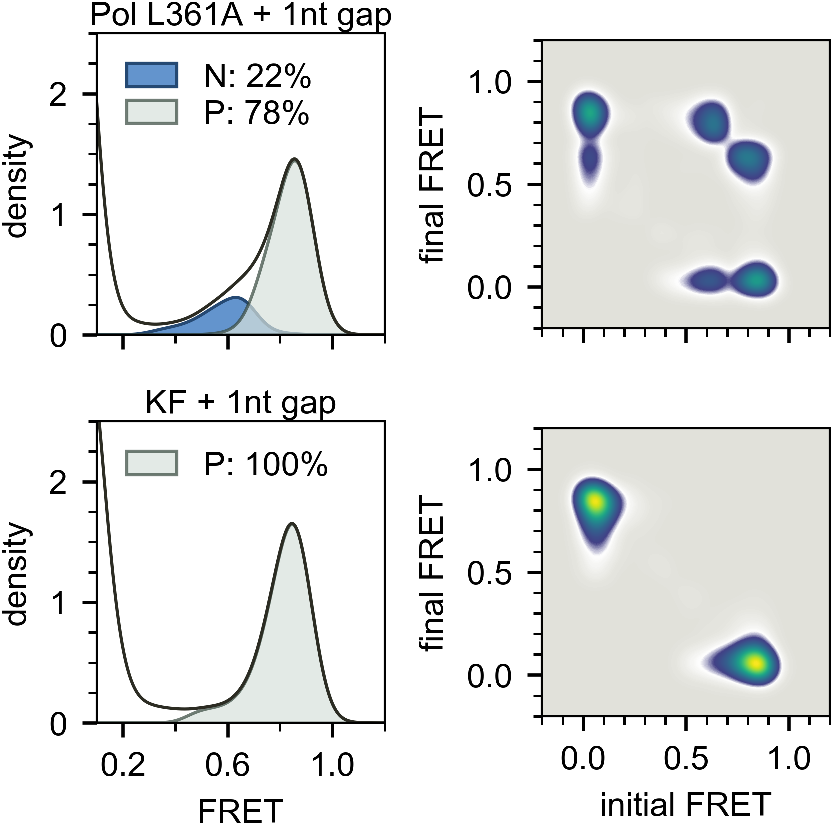
Composite FRET efficiency histograms (left) and transition density plots (right) for Pol I L361A(top panels) and KF (bottom panels) interacting with 1nt gap substrate.

**Figure 4-Figure supplement 2.**
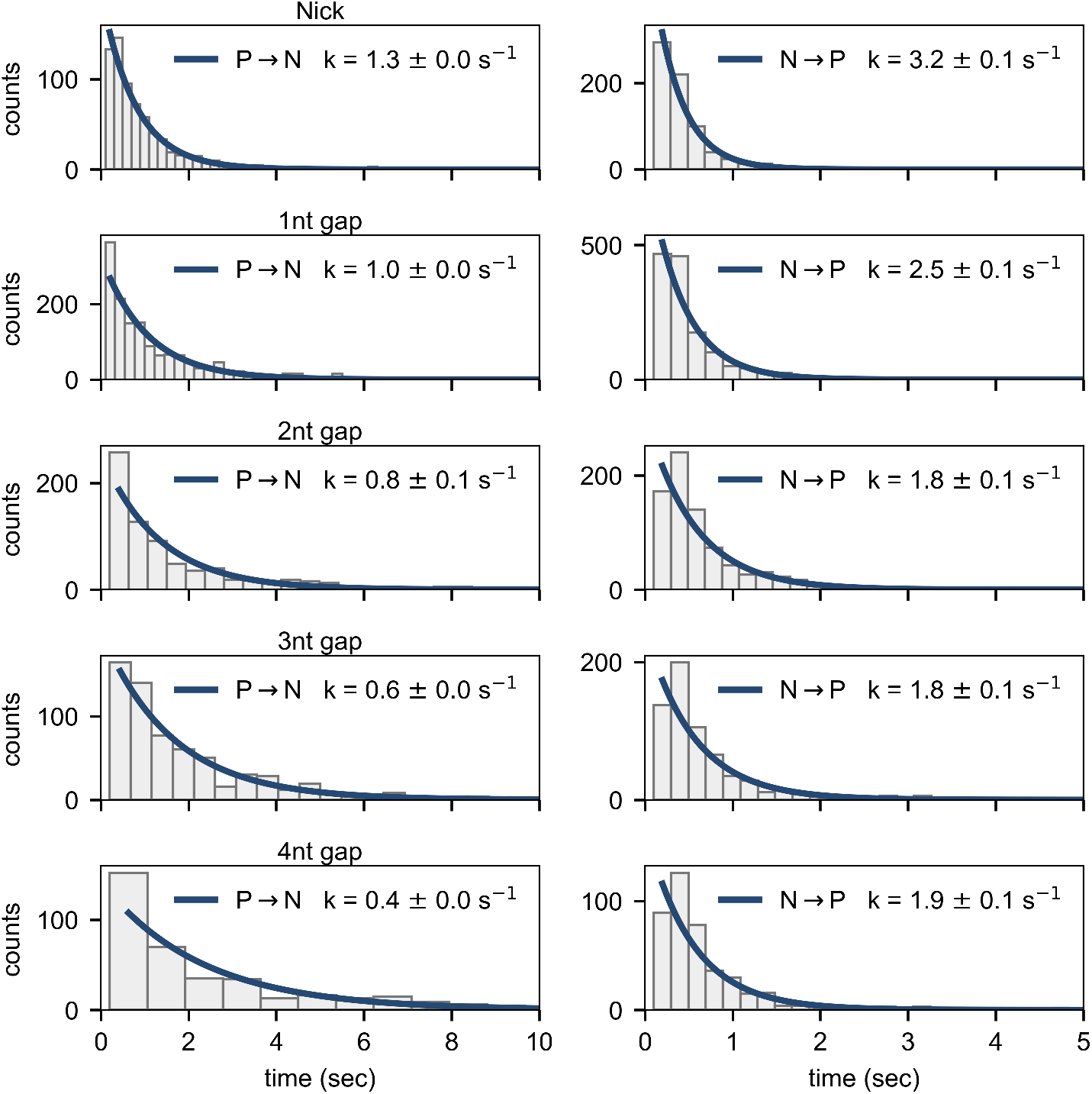
Dwell time histograms for domain-switching transitions of WT Pol I interacting with nick- or gap-containing DNA substrates. The solid lines are best fits to a single-exponential function, with the rate constants indicated.

